# Interneuron diversity and normalization specificity in a visual system

**DOI:** 10.1101/2024.04.03.587837

**Authors:** H. Sebastian Seung

## Abstract

Normalization is a fundamental operation in image processing. Convolutional nets have evolved to include a large number of normalizations (Ioffe and Szegedy 2015; Ulyanov, Vedaldi, and Lempitsky 2016; Wu and He 2018), and this architectural shift has proved essential for robust computer vision (He et al. 2015; Bjorck et al. 2018; Santurkar, Tsipras, and Ilyas 2018). Studies of biological vision, in contrast, have invoked just one or a few normalizations to model psychophysical (Mach 1868; Furman 1965; Sperling 1970) and physiological (Carandini and Heeger 2011; Shin and Adesnik 2024) observations that have accumulated for over a century. Here connectomic information (Matsliah et al. 2023) is used to argue that interneurons of the fly visual system support a large number of normalizations with unprecedented specificity. Ten interneuron types in the distal medulla (Dm) of the fly optic lobe, for example, appear to support chiefly spatial normalizations, each of which is specific to a single cell type and length scale. Another Dm type supports normalization over features as well as space. Two outlier types do not appear to support normalization at all. Interneuron types likely to be normalizers are identified not only in Dm but also in all other interneuron families of the optic lobe. For fly vision, the diversity of interneurons appears to be an inevitable consequence of the specificity of normalizations.

---

A columnar cell type (Fig. 1a) in the fly optic lobe (Fig. 1b) consists of many cells detecting the same feature at different locations in the visual field, and is analogous to a feature map in a convolutional net (Fig. 1c) (Lappalainen et al. 2023; Seung 2023). Given this close analogy, one might wonder whether fly vision also includes normalizations, which compare the activity of one neuron with the pooled activity of multiple neurons. In convolutional nets, pure spatial normalization pools over multiple locations in the same feature map (Fig. 1c, top) (Ulyanov, Vedaldi, and Lempitsky 2016). Pure feature normalization pools over multiple feature maps at the same location (Fig. 1c, bottom) (Liu et al. 2022). Spatial and feature normalization can be combined by pooling over multiple locations in multiple feature maps (LeCun, Kavukcuoglu, and Farabet 2010; Wu and He 2018). The comparison can be subtractive or divisive, and is usually a combination of both.

**Figure 1.**
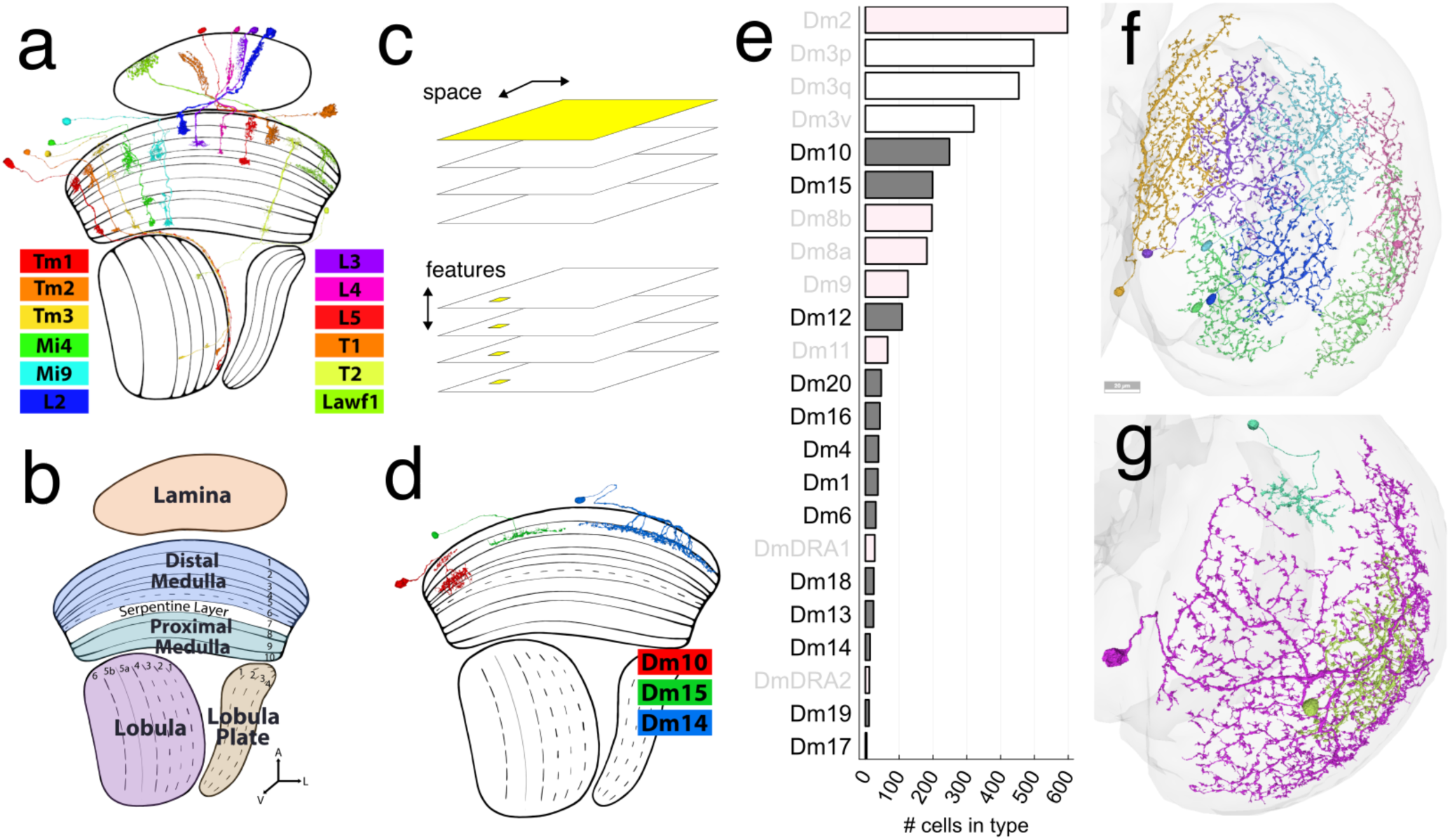
Dm interneurons and columnar neurons. (a) Some columnar cell types. Cells of each type are analogous to a feature map in a convolutional net, and are in one-to-one correspondence with ∼800 “pixels” of the fly eye. Lawf1 is an exception with a wider arbor that covers multiple pixels. (b) Main neuropils of the Drosophila optic lobe. The distal medulla is defined as the first six of ten medulla layers. (c) In convolutional nets, normalization can be purely across space, pooling over locations in the same feature map (top, yellow). Normalization can be purely across features, pooling over features at the same location (bottom, yellow). Normalization can also combine pooling across space and across features (not shown). (d) Some Dm interneuron types. Dm10 (red) is the smallest and stratifies in multiple medulla layers. Dm15 (green) and Dm14 (blue) have slightly different sizes and stratification depths, but very different synaptic partners (Figs. S1, S2). These support normalization by pooling activities of columnar cells. (e) Twenty-three Dm types ordered by the number of cells contained in each type. Dm types in dark letters are the “baker’s dozen” for which no function is known. Other Dm types (gray letters) are related to color/polarization vision (pink) or form vision (white) (Seung 2023). (f) Seven Dm6 cells. All 31 Dm6 cells would cover the entire medulla with overlap. (g) Dm15 (cyan), Dm6 (green), and Dm17 (purple) cells vary greatly in size. Cell ids 720575940609136476, 720575940634638562, 720575940636785380

A local interneuron (Fig. 1d) in the fly optic lobe pools the activity of multiple columnar neurons. Interneurons are typically inhibitory (Davis et al. 2020; Matsliah et al. 2023), and the effect of inhibition on the activity of the postsynaptic neuron is generally modeled as subtractive or divisive (Koch 2004). Therefore any cell that is downstream from an interneuron has the potential to normalize its excitatory input by subtracting or dividing the pooled activity of many columnar cells. The general idea that normalization can be implemented using local interneurons is well-known (Carandini and Heeger 2011), and has empirical support in some olfactory systems (Olsen, Bhandawat, and Wilson 2010; Zhu, Frank, and Friedrich 2013). What is new in the fly optic lobe is detailed connectomic information about the large number (>100) of interneuron types now known to exist (Matsliah et al. 2023), which makes it possible to characterize normalizations in the optic lobe with unprecedented detail.

I will start with the Dm interneuron family, which consists of 23 types found in the distal medulla (Fig. 1d, e). The functions of 13 Dm types have never been investigated (Fig. 1e).^1^ Of this baker’s dozen of Dm types, I will argue that most (ten) types support pure spatial normalizations, and are specialized for particular columnar types and length scales. For certain columnar types, spatial normalization appears to be multiscale, supported by multiple Dm types in parallel. One type (Dm13) supports feature as well as spatial normalization. The remaining two types in the Dm baker’s dozen (Dm4, Dm20) are described as interesting counterexamples that appear to not support normalization.

I conclude by searching for normalizers in all other interneuron families in the optic lobe. The long list of search results strongly suggests that normalization is a ubiquitous computation throughout the optic lobe. These findings should aid physiological studies of normalization in the fly visual system, which have recently started to emerge (Drews et al. 2020; Gür et al. 2024). More broadly, the present work addresses a question that also arises in studies of mammalian neuronal diversity: why are there so many types of inhibitory interneuron? Transcriptomic studies have identified 60 interneuron types in mouse visual cortex (Tasic et al. 2018), while functional studies differentiate between only a handful of broad inhibitory classes (Shin and Adesnik 2024). In the fly optic lobe, the reason for a large number of interneuron types may be simple. Diversity is an inevitable consequence of specificity.

## Results

### Local interneurons of the distal medulla (Dm)

Dm types have been traditionally defined by their stratification in layers of the medulla (Fig. 1d), their tangential arbor sizes (Fig. 1d, g), and their molecular properties (Fischbach and Dittrich 1989; Takemura et al. 2013; Nern, Pfeiffer, and Rubin 2015). For reliable discrimination, connectivity should also be used (Matsliah et al. 2023). The cells of each Dm type cover the medulla with overlap (Fig. 1f).

Predictions of neurotransmitter identity based on the electron microscopic images are provided by FlyWire (Eckstein et al. 2020). Predictions based on transcriptomic measurements are also available (Davis et al. 2020). Most Dm types are predicted to secrete glutamate or GABA, neurotransmitters that are usually inhibitory.

Of the twenty-three Dm types (Matsliah et al. 2023), seven (Dm8a, Dm8b, Dm9, DmDRA1, DmDRA2, Dm11, and Dm2) are known to receive direct input from photoreceptors R7 and R8, so they are presumably related to color and polarization vision (Schnaitmann, Pagni, and Reiff 2020). In a companion paper, I have provided evidence that Dm3 types (Dm3v, Dm3p, and Dm3q) are likely related to form vision (Seung 2023).

The 13 remaining Dm types (Dm1, Dm4, Dm6, Dm10, and Dm12 through Dm20) with completely unknown functions are the topic of the present work. Of this Dm baker’s dozen, ten types contain less than 100 cells each, and three types (Dm12, Dm15, and Dm10) each contain between 100 and 300 cells (Fig. 1e). In contrast, most of the Dm types with existing functional annotations contain larger numbers of cells (Fig. 1e).

### Columnar types in the distal medulla

The medulla is divided into 800 columns, which are perpendicular to its layers, and are in one-to-one correspondence with the ommatidia of the compound eye. The main synaptic partners of the Dm baker’s dozen are columnar neurons (Fig. 1a). These 12 partner types (with one exception) are confined to a single column, and each type contains about 800 cells (Fig. 1a). The activity of such a neuron mainly encodes information about the visual stimulus at or near a single ommatidium. The exception Lawf1 consists of <200 cells, each of which extends over many columns.

Lamina monopolar cells (L2, L3, L4, and L5) project from the lamina to the distal medulla. L2 and L3 receive direct input from photoreceptors in the lamina. (L1 has little or no connectivity with Dm types.) Tm1, Tm2, and Tm3 project from distal medulla to lobula. Mi4 and Mi9 project from distal to proximal medulla. T1 and Lawf1 project from distal medulla to lamina. T2 projects from distal and proximal medulla to lobula. All types are predicted cholinergic by FlyWire, except that Mi4 is predicted GABAergic, and Mi9 is predicted glutamatergic.

### Examples of spatial normalization

The shapes and orientations of the Dm arbors already hint at their functions. The arbors are typically “wide” enough to connect columnar neurons corresponding to well-separated ommatidia (Fig. 1d). Columnar types usually send information forward (from the photoreceptors towards the central brain) or sometimes backward (Lawf1 in Fig. 1a). Dm arbors look capable of sending information “sideways.” The examples of Fig. 2 will show how this lateral spread of information can enable the activity of a columnar cell to be spatially normalized relative to neighboring cells of the same type.

**Figure 2.**
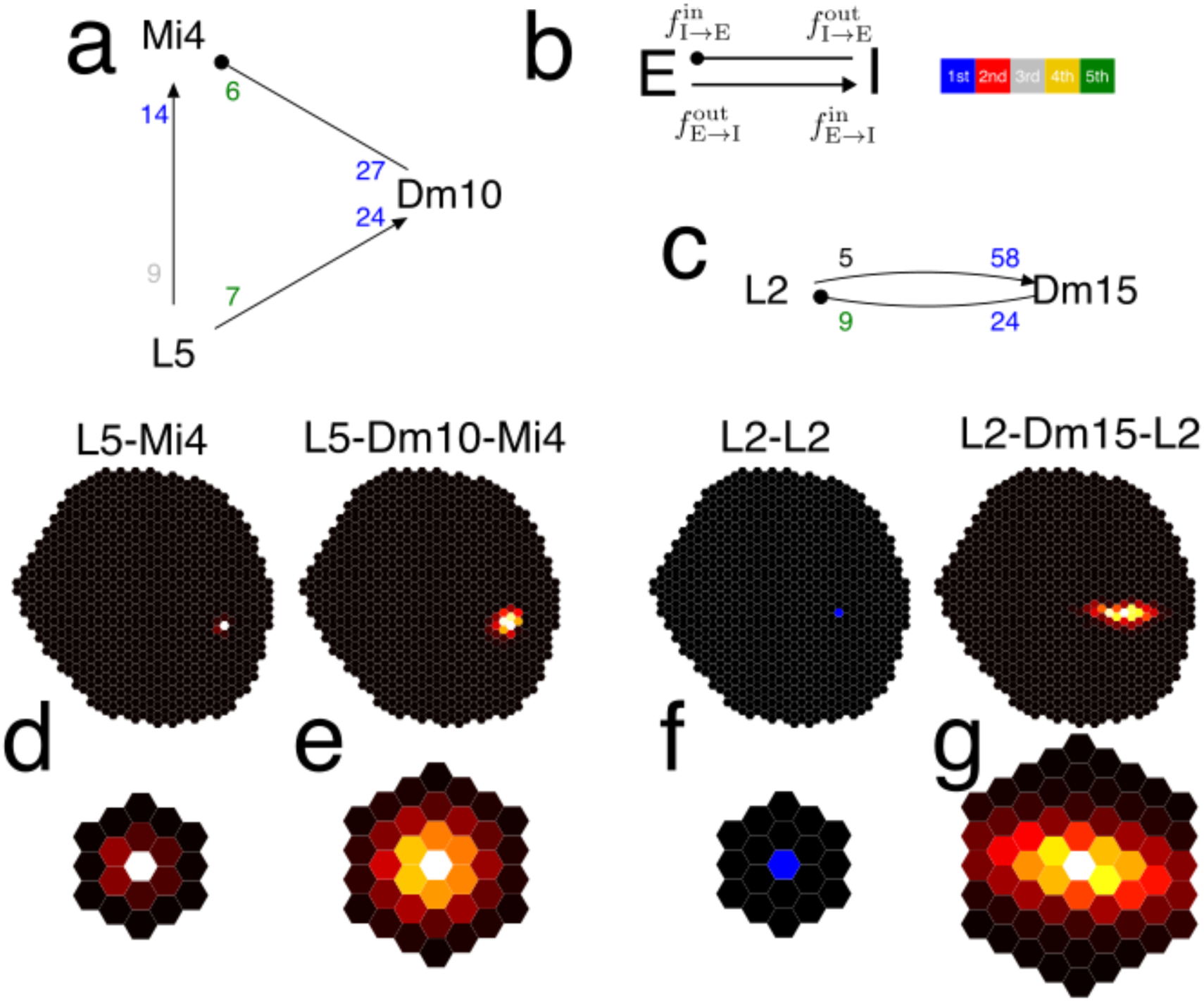
Examples of Dm-mediated spatial normalization. (a) Dm10 mediates convergent feedforward inhibition from L5 to Mi4. (b) Key to arrowheads, numbers, and colors in motif diagrams. Arrowheads indicate excitatory (triangular) and inhibitory (circular) connections. The numbers at the head and tail of an arrow are the input and output fractions (×100) of that connection. The color of the number is an ordinal ranking of strength. The input rank of a connection is obtained by sorting all connections that share the same output type in order of decreasing input fraction. The output rank of a connection is defined by sorting all connections that share the same input type in order of decreasing output fraction. (c) Dm15 mediates recurrent inhibition of L2. (d) Example and average L5-Mi4 connectivity maps. Top, all L5 cells presynaptic to an example Mi4 cell. Hexel color indicates the number of synapses from the L5 cell at that location to the Mi4 cell. Bottom, aligned and averaged for all Mi4 cells (e) Example and average L5-Dm10-Mi4 connectivity maps. Top, all L5 cells presynaptic to Dm10 cells presynaptic to the same Mi4 cell as in (d, top). Hexel color indicates the products of the L5-Dm10 and Dm10-Mi4 synapse numbers summed over all disynaptic paths from the L5 at that hexel to the example Mi4 cell. Bottom, aligned and averaged for all Mi4 cells. (f) Example and average L2-L2 connectivity maps. These maps are everywhere zero or negligible; the blue hexel just guides the eye to the location of the target L2 cell. (g) Example and average L2-Dm15-L2 connectivity maps.

Dm10 and Dm15 are the most numerous of the types in the Dm baker’s dozen (Fig. 1e). Clues to their functions can be gleaned from their presynaptic and partner types. L5 is the strongest input to Dm10 (Figs. 2a, S1), and Mi4 is the strongest output of Dm10 (Figs. 2a, S2). Connection strengths are quantified in Fig. 2a following the key in Fig. 2b.

The indirect pathway L5-Dm10-Mi4 is paralleled by a direct L5-Mi4 connection (Fig. 2a). L5 is cholinergic and presumed excitatory, while Dm10 is predicted GABAergic and presumed inhibitory. It follows that the direct pathway is excitatory, while the indirect pathway is net inhibitory. Since both direct and indirect pathways converge onto the same target, the Dm10 circuit motif (Fig. 2a) will be called “convergent feedforward inhibition,” or “feedforward inhibition” for short.

For Dm10 the strongest input and output were different columnar types. For Dm15, it turns out that the strongest input and output are the same columnar type, L2 (Fig. 2c, S1, S2). Dm15 is predicted glutamatergic and presumed inhibitory. The motif of Fig. 2c will be called “recurrent inhibition,” meaning that L2 inhibits itself through Dm15.

It might seem counterproductive for L5 to exert opposite effects on Mi4 through the direct and indirect pathways in Fig. 2a. But this diagram ignores spatial organization. Through the direct pathway, Mi4 is excited by L5 in the same column, with negligible input from L5 in neighboring columns (Fig. 2d). Tracing all indirect pathways from L5 cells to Dm10 cells to an Mi4 cell yields an estimate of the L5-Dm10-Mi4 “inhibitory field” (Fig. 2e). This is an estimate of how much the activity of an Mi4 cell is inhibited by L5 activity in neighboring columns. Figs. 2d and 2e mean that an Mi4 cell compares its direct input from the L5 cell in the same column with Dm10-pooled activity of L5 cells in neighboring columns. Mi4 activity is driven not by absolute L5 activity, but L5 activity relative to other L5 cells, so this is an example of spatial normalization.

A similar analysis predicts the L2-Dm15-L2 inhibitory field (Fig. 2g). Here also the normalization is spatial, because it enables the L2 cell to compare its own activity with that of other L2 cells.

### Dm cells are dominated by one or two input types

The above analysis raises two questions. First, the Dm10 and Dm15 circuit motifs (Fig. 2a, c) depicted only the strongest input and output to these Dm types. Is it really a good approximation to neglect the others? Second, does the analysis extend from Dm10 and Dm15 to other Dm types?

These questions can be addressed by examining the bar graphs of Fig. S1, which show the top four input fractions for all Dm types. The strongest input to a Dm type (the leftmost bar in the graphs) will be called the “winner.” For each Dm type, the winner is always some type of lamina monopolar cell (L2 through L5, Fig. S1), except that Tm2 is the winner for Dm16. The second strongest input to a Dm type will be called the “runner-up.” For most Dm types, the winner is much larger than the runner-up and all other inputs. The exceptions are Dm13, Dm4, and Dm20, for which the runner-up is nearly tied with the winner. Therefore the following rule holds:

> Rule 1. Most Dm types in the baker’s dozen are dominated by a single input type.

This notion of dominance can be formalized by ordering the inputs (as in Fig. S1), and then quantifying the fractional decrease of each successive input. Find the smallest *i* such that input *i*+1 is less than γ times input *i*. Then we say that inputs 1 through *i* are dominant. Setting γ=0.6, we see that either one or two inputs are dominant for each type. These are the inputs above the red line in Fig. S1, the height of which is set by γ times input *i*. The choice of threshold parameter γ is somewhat arbitrary, but modest changes do not qualitatively change the overall picture of high input specificity.

### Dominant outputs of Dm types

The output fractions of each Dm type are shown in Fig. S2. Seven types are dominated by a single output, and three types are dominated by two outputs. A few types are dominated by large output sets (Dm1 and Dm6 by four outputs, and Dm19 by five outputs). The top outputs of the Dm cells are more diverse than the top inputs (Fig. S2). They include all the columnar types that are top input of some Dm cell (L2, L3, L5, Tm2) except for L4. In addition, the top outputs include Tm1, Mi1, Mi4, T2, and Lawf1. All of the top outputs are confined to single columns, except for Lawf1 which extends over multiple columns. For five Dm types, the top output is the same as the top input.

### Dm types supporting chiefly spatial normalization

Fig. 3 depicts the connectivity of Dm types in the baker’s dozen that have a single dominant input. These Dm types perform chiefly spatial pooling; they do not pool over features to a first approximation. The following simple rule about their postsynaptic targets almost always holds:

> Rule 2. An output target of a Dm type is either the same as its dominant input source (recurrent inhibition), or cell types that are directly downstream of its dominant input source (feedforward inhibition).

**Figure 3.**
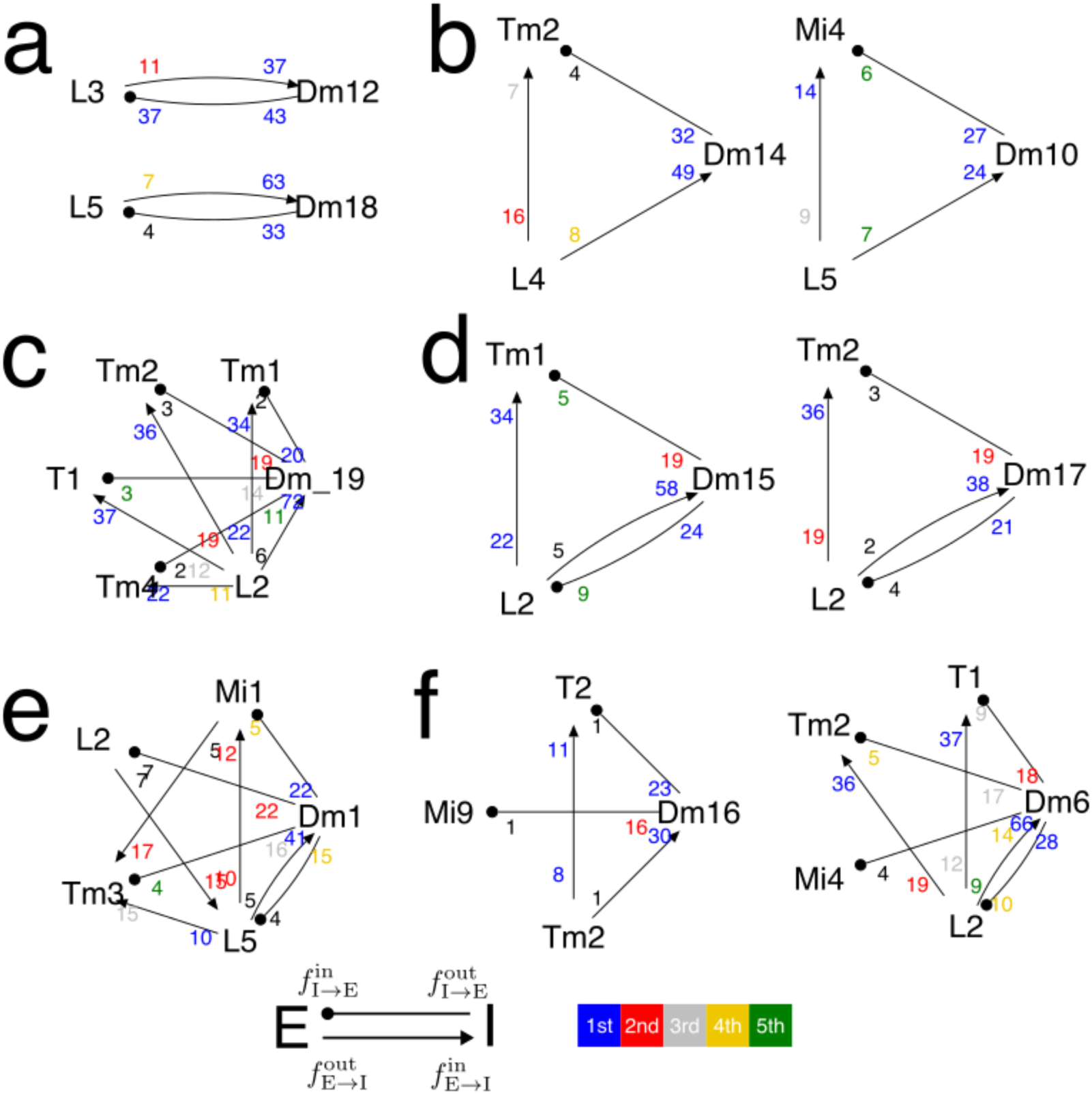
Spatial normalization is common. Dm types with their dominant inputs and outputs (defined in Fig. S1, S2). Only connections with importance (Methods) exceeding a 10% threshold are shown. Motifs largely obey Rule 2, meaning that normalization is purely spatial. (a) Dm12 and Dm18 mediate recurrent inhibition of L3 and L5, respectively. (b) Dm14 and Dm10 mediate feedforward inhibition from L4 and L5, respectively. (c) Dm19 mediates feedforward inhibition from L2 to multiple targets. (d) Dm15 and Dm17 mediate a combination of feedforward and recurrent inhibition. (e) Dm1 conforms to Rule 2, and also contains “extra” connections between two target types (Mi1 to Tm3) and from target to source type (L2 to L5). (f) Dm16 and Dm6 conform to Rule 2, except for the “dangling” connection from Dm6 to Mi4. The connection from Dm16 to Mi9 is only apparently dangling; a connection from Tm2 to Mi9 does exist but is below the threshold for visibility in the diagram.

This promotes the specific example motifs in Figs. 2a and 2b to be a general rule for the Dm baker’s dozen. Rule 2 means that the activity of a columnar cell is compared with the pooled activity of cells of the same type. In other words, the normalization is chiefly spatial.

To illustrate Rule 2, Fig. 3 shows each Dm type along with its dominant input and output types. For clarity, only connections with input fraction or output fraction greater than 10% are included; weaker connections are omitted from the diagrams.

Dm12 (Takemura et al. 2015; Xu et al. 2018) and Dm18 are reciprocally connected with L3 and L5, respectively (Fig. 3a), and therefore conform to recurrent inhibition, similar to Fig. 2b.

Dm14 conforms to feedforward inhibition (Fig. 3b), just like the Dm10 motif that was in Fig. 2a and is repeated in Fig. 3b for convenience. The Dm19 motif looks more complex (Fig. 3c), but it is simply four feedforward inhibition motifs that overlap.

Dm15 and Dm17 combine recurrent and feedforward inhibition (Fig. 3d). For example, one target of Dm15 (Tm1) is feedforward inhibition, while the other target (L2) is the recurrent inhibition that was shown in Fig. 2b.

The Dm1 diagram (Fig. 3f) fully conforms to Rule 2, though there are also “extra” connections.

As combinations of recurrent and feedforward inhibition, the Dm16 and Dm6 diagrams (Fig. 3e) largely obey Rule 2, but also contain “dangling” outputs Mi9 and Mi4, which do not conform. It turns out that the Dm16 diagram does not truly lack a connection from Tm2 to Mi9; there is a weak connection that does not show up because it is below the chosen threshold for being displayed. So Dm16 actually does conform to Rule 2.

On the other hand, the Mi4 target of Dm6 really does violate Rule 2, because the direct connection from L2 to Mi4 is vanishingly small. With a broader definition of normalization, one could say that Mi4 is normalized by pooled L2 activity. A stricter definition of normalization would require that the target also be an input to the pooling. (This definition would be analogous to normalization of a probability distribution, for which the numerator occurs in the sum in the denominator.) Rather than normalization, the L2-Dm6-Mi4 pathway could be interpreted as mediating an opponent interaction between L2 and Mi4. This is a case of ON-OFF opponency, because L2 is OFF while Mi4 is ON.

Overall, each Dm type tends to devote large fractions (>10%) of its input and output synapses to connections that conform to either recurrent or feedforward inhibition. In this sense, it seems apt to say that the Dm type “serves” or “supports” its dominant input. On the other hand, the dominant input usually devotes a small fraction (<10%) of its output synapses to the Dm type, and the targets of the dominant input usually receive a small fraction (<10%) of its input synapses from the Dm type. In this respect, the relationship between the Dm type and the columnar types is asymmetrical.

### Spatial normalization is multiscale

Figure 3 depicted connectivity from the viewpoint of each Dm type. It is also helpful to visualize connectivity from the viewpoints of columnar types. Three Dm types provide recurrent inhibition to L2 (Fig. 4a), and two Dm types provide recurrent inhibition to L5 (Fig. 4b). This might seem redundant, but visual examination of the types reveals that their properties are quite different. For the Dm partners of L2, the number of cells per type varies from 200 to 30 to 4 (Dm15, Dm6, Dm17, Fig. 1c). The arbors of these types likewise range from small to medium to large (Fig. 4g). Dm6 is predicted GABAergic, Dm15 is predicted glutamatergic, and Dm17 has no confident prediction.

**Figure 4.**
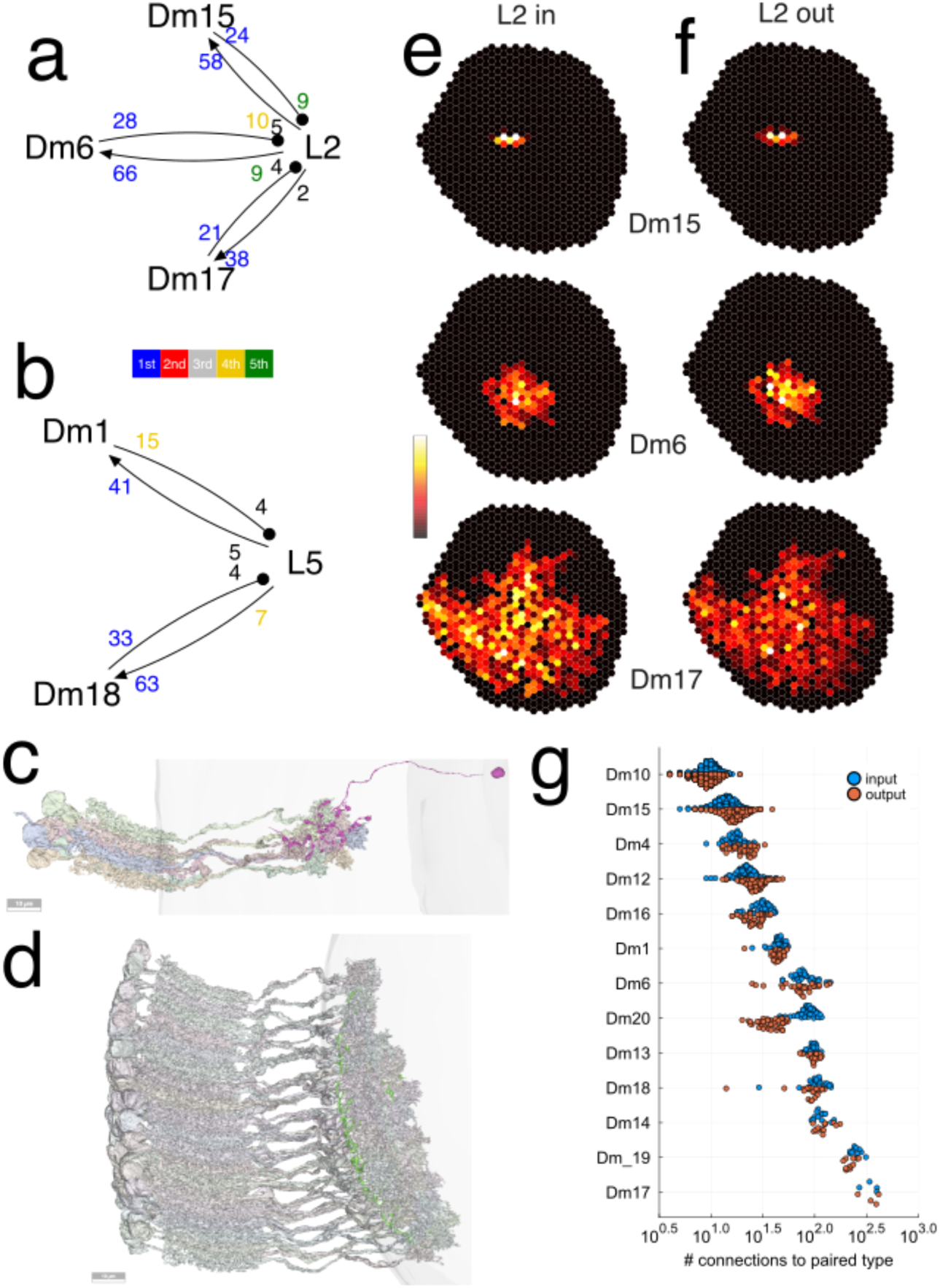
Spatial normalization is multiscale. (a) Recurrent inhibition of L2 is mediated by three Dm types. (b) Recurrent inhibition of L5 is mediated by two Dm types. (c) Dm15 cell (red, 720575940644990062) and 9 presynaptic L2 cells (>4 synapses) (d) Dm6 cell (green, 720575940609136476) and 74 presynaptic L2 cells (>4 synapses) (e) L2-Dm connectivity maps for example Dm15, Dm6, and Dm17 cells. (f) Dm-L2 connectivity maps for example Dm15, Dm6, and Dm17 cells. The inputs and outputs to a Dm cell have the same spatial organization, because they correspond to the columns covered by the Dm arbor. Colorbar maximum 12, 37, 15 synapses for L2-Dm and 16, 18, 17 synapses for Dm-L2. Cell IDs are 720575940609136476, 720575940634638562, 720575940644990062. (g) For cells of each Dm type, number of connections with strongest input and output types. For each Dm type, the distributions are similar for input and output. Dm20 is an exception because its strongest output is Lawf1, which is multicolumnar.

L2 cells extend from the lamina to the medulla, where they synapse onto Dm15 (Fig. 4c), Dm6 (Fig. 4d), and Dm17 cells. Example maps of L2-Dm connectivity are shown in Fig. 4e, and indicate the pool of L2 cells over which a Dm cell pools. The maps can also be regarded as predictions of Dm receptive fields. Example maps of Dm-L2 connectivity are shown in Fig. 4f. Through a Dm cell, an L2 cell on one side of the Dm arbor can indirectly inhibit an L2 cell on the other side of the arbor. The L2-Dm15-L2 inhibitory field (Fig. 2h) is wider than the L2-Dm15 (Fig. 4e, top) and Dm15-L2 (Fig. 4f, top) maps, because the inhibitory field of an L2 cell is mediated by the multiple Dm15 cells that are presynaptic to it. L2-Dm6-L2 and L2-Dm17-L2 inhibitory fields (data not shown) are wider than the L2-Dm15-L2 inhibitory field (Fig. 2h), because the range of lateral inhibition is set by the spatial extent of the Dm arbor.

To summarize, L2 appears to be simultaneously normalized by three Dm types that likely operate at different length scales. The example cells (Fig. 4e, f) illustrate the range of variation across types. Similarly, L5 is normalized by two Dm types that likely operate at different length scales (Fig. 4b). A more complete picture of intra-type as well as inter-type variation in Dm arbor size is provided by Fig. 4g.

### Outlier Dm types

Of the Dm baker’s dozen, three types (Dm13, Dm4, and Dm20) are omitted from Fig. 3 because they are dominated by two input types (Fig. S1). Dm13 pools Tm2 and L5 activity over space, and this signal is used to normalize Tm2 activity (Fig. 5a). Therefore, Dm13 supports feature normalization (Fig. 1c, bottom) as well as spatial normalization, and deviates somewhat from the pure spatial normalizers of Fig. 3.

**Figure 5.**
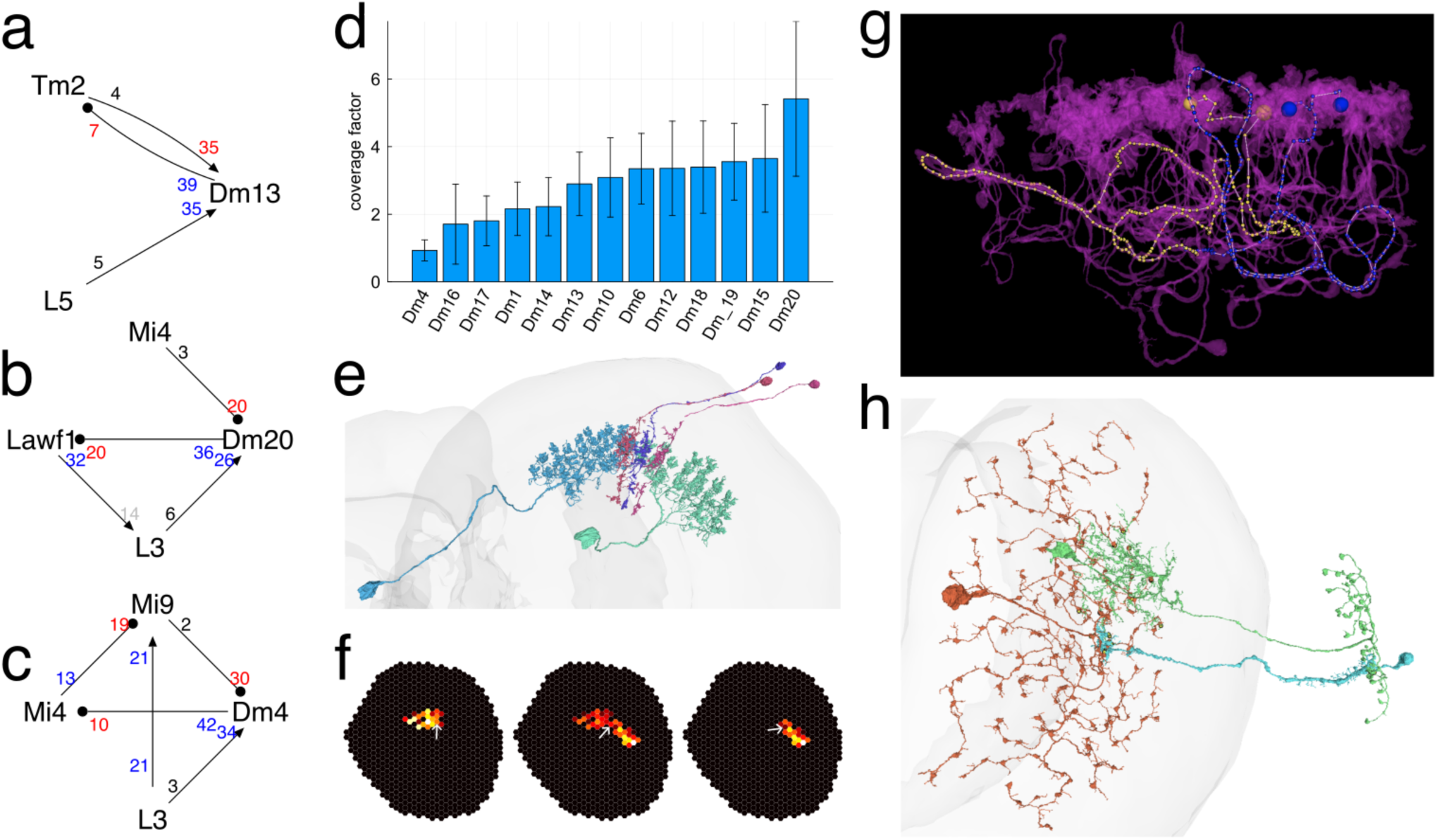
Exceptions to chiefly spatial normalization. (a) Dm13 supports feature normalization, pooling activity over both Tm2 and L5, as well as spatial normalization. (b) Dm20 does not appear to support normalization, as it receives inhibitory input from Mi4. (c) Dm4 does not appear to support normalization, as it receives inhibitory input from Mi9. (d) Coverage factors of Dm types. Dm4 is the only type with coverage factor close to one, meaning almost perfect tiling of the medulla (Fig. S4). Dm20 has an unusually large coverage factor. (e) Mi4 receiving input from two Dm4 cells, flanked by two Mi4 receiving input from single Dm4 cell. Cell ids are 720575940613047794 720575940613604581,720575940643591368 720575940620113969 720575940614309416. (f) Example L3-Dm4-Mi4 connectivity maps for the Mi4 cells of (b). The white arrow in each map points to the location of the Mi4 cell. The dramatic differences between these maps constitute a severe breaking of translation invariance. (g) Dm4 may not be electrotonically compact, because of long path lengths between columns. Two neighboring columns (large blue and large yellow balls indicate two pairs of terminals in the two columns) are separated by a Euclidean distance of roughly 7 µm, but the neurite path length (blue and yellow skeletons) between the columns exceeds 200 µm. The blue path from the blue terminals and the yellow path from the yellow terminals meet at a branch point. Each path exceeds 100 µm. Cell id is 720575940627092016 in v630. (h) Example of L3 (cyan) to Dm20 (orange) to Lawf1 (green) mediating 3-cycle recurrent inhibition. Lawf1 is a centrifugal cell that projects from the medulla to the lamina (left to right), where it synapses onto L3. Cell ids are 720575940607554434 720575940608960009 720575940633352100.

On the other hand, Dm20 (Fig. 5b) and Dm4 (Fig. 5c) appear to deviate fundamentally from the conception of normalization, because they receive inhibitory columnar inputs Mi4 (predicted GABAergic) and Mi9 (predicted glutamatergic), respectively. By definition, the pool for normalization should contain excitatory neurons only. A neuron that receives both excitatory and inhibitory columnar inputs does not fit the notion of pooling for normalization, in the same way that normalizing a probability distribution does not extend to numbers of mixed sign.

Although Dm4 and Dm20 are outliers, they are worth discussing in more detail because they are the exceptions that prove the rule. They differ from the spatial normalizers in Fig. 3 in multiple ways that are illuminating. They violate Rule 2 as well as Rule 1, and they are also unusual because of their spatial coverage.

Each Dm type covers the medulla, in the sense that every column is covered by at least one cell of that type. In vertebrate retina, the coverage factor of a neuron type has been defined as the average number of neurons of that type covering a single point in the retina, and has been estimated by physiological and anatomical methods (Sterling 1983). Here coverage factor can be quantified for the first time using anatomical connectivity. For a given Dm type, choose a columnar type that is a synaptic partner. For all cells of the columnar type, count the number of partner cells of the Dm type, and then average over columnar cells. This is done in Fig. 5d using the strongest input type for each Dm type. Most of the Dm baker’s dozen have coverage factors close to two or three.

Dm4 and Dm20 are outliers in the graph. Dm4 has the lowest coverage factor, which is close to one (Fig. 5d). This is consistent with visual inspection, which reveals that Dm4 cells overlap very little, fitting together beautifully, almost like floor tiles (Fig. S4). Dm20 has the highest coverage factor, which exceeds five (Fig. 5d).

### Overlap supports translation invariance

In vertebrate retina, it has often been noted that coverage factor of cell types is typically greater than one, at least for amacrine and ganglion cells (Bae et al. 2018). I claim that overlapping coverage plays a simple role for visual function, which is to support translation invariance. The converse of this assertion, that translation invariance breaks down when there is no overlap, is easy to illustrate for Dm4. If lateral inhibition were the function of Dm4, then the almost perfect tiling of Dm4 would have marked effects near the borders of Dm4 arbors. This is illustrated by three adjacent Mi4 cells in Fig. 5e. The middle Mi4 cell receives input from two Dm4 cells, while the flanking Mi4 cells each receive input from a single Dm4 cell. Fig. 5f shows the consequences for L3-Dm4-Mi4 connectivity maps. The middle Mi4 cell receives disynaptic input from L3 cells on its left and right. The left Mi4 cell receives input from L3 cells on its left, and the right Mi4 cell receives input from L3 cells on its right. Such marked differences in the inhibitory fields of adjacent Mi4 cells would be observable through visual physiology, and would contradict the concept of translation invariance, that cells of the same type perform the same computation at different locations in the visual field.

### Dm4 likely does not pool over space

However, there is an alternative scenario. Perhaps Dm4 does not mediate lateral inhibition after all. Zooming into the arbor of a Dm4 cell reveals that its neurites take amazingly circuitous paths to travel from one column to the next (Fig. 5g). The neurite path length between two neighboring columns turns out to exceed 200 µm, even though the columns are separated by a Euclidean distance of just 7 µm. This is because of the looping and auto-fasciculation of the Dm4 arbor. This is not the case for the arbors of other Dm types, in which neurites take straighter paths.

I predict that the long looping neurites serve to electrically isolate columns from each other. Each column of the Dm4 arbor is predicted to be an independent functional unit. If this is the case, Dm4 mediates unicolumnar interactions, and the connectivity maps of Fig. 5f are not actually lateral inhibitory fields. Such an “extreme compartmentalization” has been observed in the CT1 amacrine cell of the optic lobe (Meier and Borst 2019).

If Dm4 is electrically compartmentalized, it is natural to wonder whether the same could be true of the other Dm types. The claim that the Dm types of Fig. 3 are spatial normalizers implicitly assumed that these cells are electrotonically compact, meaning that synaptic input causes an electrical effect that spreads across the entire cell with little or no attenuation. This seems plausible for the small Dm10 and Dm15 cells of Fig. 2, but is less clear for large Dm cells. It is certainly conceivable that the large Dm types do not spatially pool over the entire extent of their arbors. But Dm4 is an outlier in the baker’s dozen; the other Dm types do not have the same circuitous neurites. Due to this stark contrast, the starting assumption that the other Dm types do pool over space seems reasonable unless evidence to the contrary emerges.

### Dm20 mediates feedback lateral inhibition

The Dm20 graph (Fig. 5b) contains a 3-cycle L3→Dm20→Lawf1→L3. Lawf1 is predicted cholinergic, consistent with transcriptomic evidence (Davis et al. 2020), though immunolabeling suggests that Lawf1 is both cholinergic and glutamatergic (Yuan et al. 2021). Assuming that Lawf1 excites L3, and Dm20 inhibits Lawf1, the 3-cycle is net inhibitory. Three cells in such a 3-cycle are shown in Fig. 5h.

Lawf1 is a columnar neuron, by the traditional definition (Fischbach and Dittrich 1989), because its axon is oriented parallel to the main axis of the columns. However, it has two unusual properties. First, Lawf1 dendritic and axonal arbors cover many columns. The other columnar partners of Dm cells are mostly confined to a single column. This difference is related to the fact that Lawf1 is much less numerous than the other columnar types (Fig. 1a). Second, Lawf1 is a centrifugal neuron, projecting from the medulla to the lamina. In summary, Dm20 likely mediates lateral inhibition between L3 neurons, but the lateral inhibition involves feedback to the lamina.

### Dm-Dm disinhibition

Up to now, only connectivity of Dm types with columnar types has been discussed. Dm-Dm connectivity is shown in Fig. S5a. Pixel values indicate the “importance” of the connection, defined as the maximum of input and output fraction (Methods). Some important inter-type Dm connections exist (Fig. S5a), though connectivity is rather sparse. Thresholding importance at 5%, there are only 12 out of 156 possible connections between distinct Dm types. Dm13 and Dm17 have no connections that survive thresholding. The thresholded graph of important connections is acyclic/feedforward (Fig. S5b), the only exception being that Dm10 and Dm20 mutually inhibit each other.

It is not obvious how Dm-Dm connectivity fits into the conceptual framework of normalization. One interesting case is Dm19 inhibition of Dm6 (Fig. S5a). Since both types share L2 as source (Figs. 3c, e), Dm19 can be interpreted as normalizing L2 input to Dm6. In other words, this could be a case of normalizing a normalizer.

All connections above the diagonal in Fig. S5a are from larger to smaller types. Some connections also exist below the diagonal, from smaller to larger types. Intra-type connectivity is weak, with input and output fractions less than 1% for 10 out of 13 types (diagonal pixels, Fig. S5a).

### Normalization in other interneuron families

We have seen that most Dm interneurons in the baker’s dozen are likely to act as normalizers. Does this finding generalize to interneuron families other than Dm? Space does not permit a full exploration of this question, but it is straightforward to identify candidate normalizers by searching for strong recurrent loops like Fig. 2b. It turns out that such loops exist in all other interneuron families (Pm, Sm, Li, and LPi) of the optic lobe that were described by (Matsliah et al. 2023).

Candidate Pm normalizers (Fig. 6a) interact with columnar cell types in the ON (Mi1) and OFF (Tm1, Tm2) channels that were defined in (Matsliah et al. 2023). Notably Pm08 interacts with both Mi1 and Tm1, so it normalizes over features (ON and OFF) as well as space.

**Figure 6.**
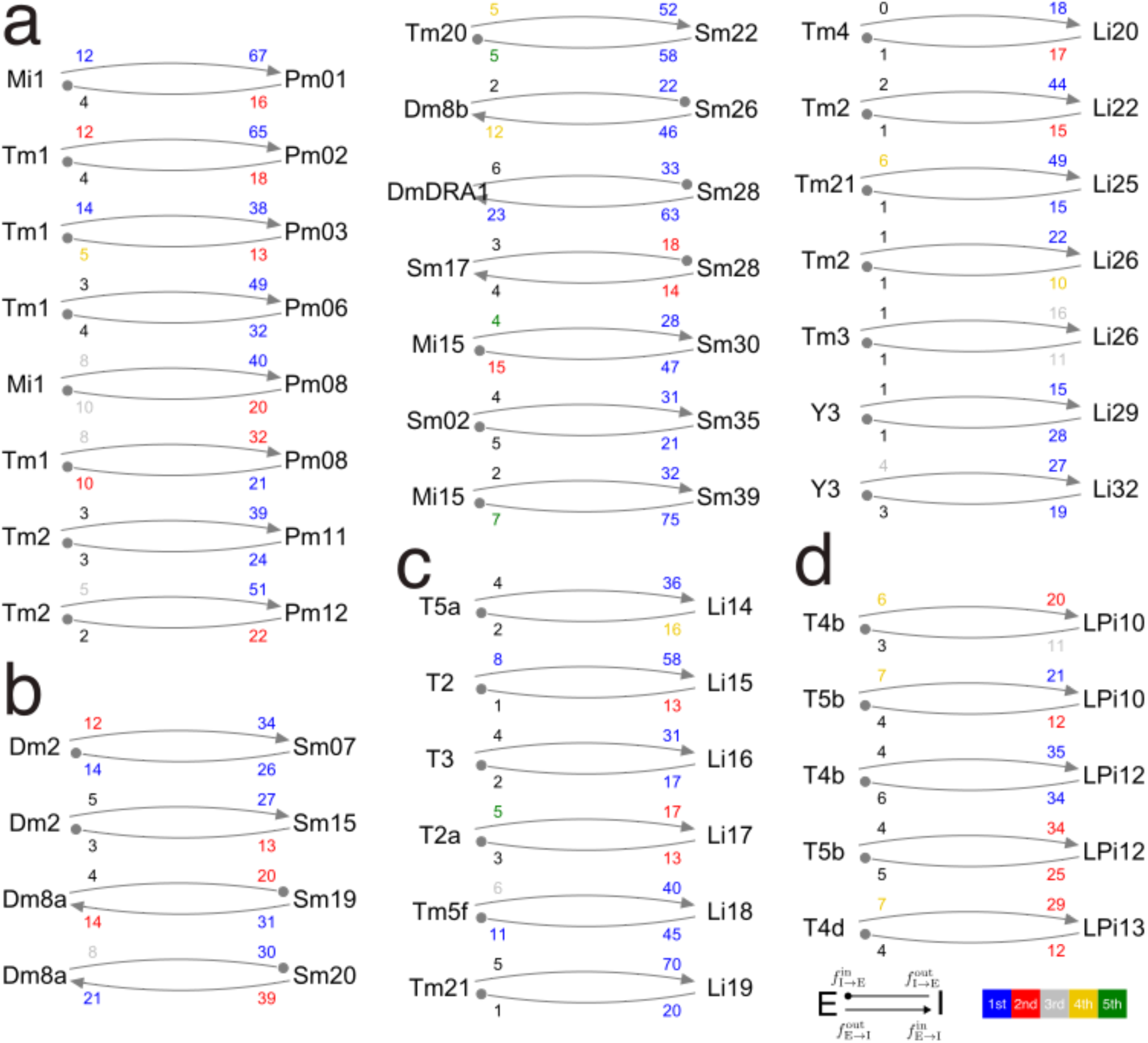
Candidate interneuron normalizers. In all motifs, left type is normalizee and right type is normalizer. Normalizee is source of more than 10% of input synapses and target of more than 10% of output synapses of normalizer (percentages are small numbers near the normalizer). In almost all cases, normalizee is the strongest or second strongest input and output of normalizer (almost all percentages are blue or red). Normalizers include interneuron types in the Pm (a), Sm (b), Li (c), and LPi (d) families. Some normalizees are connected to multiple normalizers, similar to Fig. 4, and some normalizers are connected to multiple normalizees.

Candidate Sm normalizers (Fig. 6b) interact with targets (Dm2, Dm8a, Dm8b, Tm20, DmDRA1, Mi15) of inner photoreceptor synapses (Kind et al. 2021), as well as with two Sm types. In several cases, the normalizer is excitatory while the normalizee is inhibitory. More analysis is required to determine whether these motifs can be interpreted as normalizing in spite of the sign inversion.

Candidate Li normalizers (Fig. 6c) interact with types related to object (T2, T2a, T3, Tm21, Y3), motion (T5a), and color (Tm5f) vision, as well as types in ON (Tm3) and OFF (Tm2, Tm4) channels.

Candidate LPi normalizers (Fig. 6d) interact with T4/T5 types. These motifs suggest that normalization can be a function of LPi types. Previously, the only proposed function of LPi types was motion opponency (Mauss et al. 2015; Ammer et al. 2023).

Some columnar types are connected to multiple candidate normalizers in the same interneuron family. As in Fig. 4, this could implement normalization at multiple length scales. Furthermore, a single columnar type can be normalized by interneurons distributed across multiple neuropils (Figs. 3, 6). For example, Tm2 is likely normalized by Dm, Pm, and Li types, a rich and complex array of overlapping mechanisms.

All told, Fig. 6 contains 32 interneuron types that are candidate normalizers. The length of the list depends on a threshold, which was set at 10% of the input and output fraction of the normalizer (Fig. 6 caption). The list would become longer if the threshold were lowered, or if convergent feedforward inhibition like Fig. 2a were included. The list could shorten if types turn out to be electrically compartmentalized, as suspected for Dm4. Therefore Fig. 6 is not intended to be exhaustive; it is merely meant to support the overall claim that a significant fraction of interneuron types throughout the optic lobe are likely to be normalizers.

## Discussion

A connectomic analysis suggests that the fly distal medulla contains many highly specific normalizations. According to Rule 1, most Dm types in the baker’s dozen pool activity over a single source type at multiple locations (Fig. S1). According to Rule 2, the target type is either the same as the source type (recurrent inhibition), or the target type is directly downstream from the source type (convergent feedforward inhibition). A Dm interneuron type can mediate either recurrent or feedforward inhibition, or a combination of both (Fig. 3).

Normalization was classically assumed to have one characteristic length scale, but multiple scales turn out to exist in the fly distal medulla. For example, the notion that “the activity of an L2 cell is normalized by the pooled activity of other L2 cells,” even if specific to the L2 cell type, is still coarser than reality. L2 appears normalized by three Dm types working at different spatial scales (Fig. 4a).

Edge cases and outliers are useful for clarifying the definition of normalization. If one requires that the normalization pool contain the target cell type, then the lateral inhibition received by Mi4 from L2 via Dm6 (Fig. 3f) is not normalizing but can be interpreted as ON-OFF opponency. On the other hand, the lateral inhibition received by Tm2 from L5 via Dm13 is normalizing because Dm13 pools over both Tm2 and L5. Dm13 is the one type in the baker’s dozen that supports feature as well as spatial normalization. If one requires that the normalization pool contain only excitatory neurons, then Dm20 and Dm4 are likely not normalizers because their inputs from Mi4 and Mi9 are predicted inhibitory.

### Differences with convolutional nets

The motifs of Fig. 3 are analogous to spatial normalizations in a convolutional net, but there are some important differences. The first is that Dm types pool over spatial neighborhoods. In convolutional nets, the most popular pure spatial normalizations pool over all of visual space (Ulyanov, Vedaldi, and Lempitsky 2016). Spatial normalization over neighborhoods is generally used in combination with feature normalization in convolutional nets (Fukushima 1975, 1980; LeCun, Kavukcuoglu, and Farabet 2010). Spatial normalization by Dm types can be multiscale (Fig. 4), which is not common in convolutional nets. The combined spatial and feature normalization of Dm13 (Fig. 5a) is similar to group normalization (Wu and He 2018), except that one of the members of the group (L5) is targeted by Dm13 only very weakly.

The recurrent inhibition motif (Figs. 2c, 3a) is analogous to normalizing a feature map in a convolutional net, because it normalizes a single cell type, and its effects are seen by all downstream cell types. Convergent feedforward inhibition (Fig. 2a, 3b) is potentially more specific. In this motif, a target type compares source activity relative to some local average of source activity. A different target of the same source could compute a different normalization of the source or even none at all. In a convolutional net, this would be a different kind of normalization associated not with a feature map but with a connection between feature maps.

### Dm4 is predicted to be electrically compartmentalized

The attenuation of electrical signals along a passive neurite has been studied theoretically (Koch 2004). Such theories cannot generate accurate quantitative predictions about Dm cells because of uncertainty in parameters like the specific transmembrane resistance and the axoplasmic resistivity. These parameters may vary by a factor of two or more, even after constraining them with electrophysiological measurements (Gouwens and Wilson 2009).

Since it is difficult to predict the absolute degree of Dm cell compartmentalization, I have resorted to comparing Dm cell morphologies to predict the relative degree of compartmentalization. Unique among the Dm baker’s dozen, the Dm4 cell has long looping neurites that make the path length between neighboring columns many times greater than the Euclidean distance. This outlier morphology suggests that Dm4 is much more electrically compartmentalized than other Dm types.

Similar long looping neurites have been observed in the CT1 cell, an example of a large amacrine cell in the optic lobe that exhibits “extreme compartmentalization.”(Meier and Borst 2019) Because of these long neurite paths, CT1 terminals in the medulla are electrically isolated from each other to a first approximation, even those in neighboring columns. Calcium imaging experiments show that CT1 terminals act as independent functional units; each CT1 terminal responds to visual input in its own column (Meier and Borst 2019). Electrical compartmentalization has also been reported in interneurons of the antennal lobe (Taisz et al. 2023) and mushroom body (Amin et al. 2020) of the olfactory system.

Cable theory predicts that attenuation depends more strongly on path length, and rather weakly on diameter and membrane properties (due to a square root in the formula for length constant) (Koch 2004). Therefore varying path length across Dm types should be an effective way of varying degree of compartmentalization.

### Translation invariance

Convolutional nets are often said to be translation invariant, meaning that the neurons of a feature map perform the same computation at different locations in the visual field. However, it is well-known that translation invariance is broken if feature maps are downsampled. For example, some convolutional nets contain an operation known as strided convolution that subsamples the result of regular convolution (Dumoulin and Visin 2016). A stride of two means that every other pixel is retained, resulting in a smaller output feature map with a spatial resolution that is halved relative to the input feature map.

The stride of a convolution controls coverage. If the stride is equal to the kernel size, then the coverage factor is exactly one, similar to Dm4. If the stride is equal to one, the coverage factor is equal to the kernel size, and translation invariance holds. A stride greater than one means that translation invariance is broken, with more severe effects for larger stride.

Each Dm type is analogous to a feature map that is constructed via strided convolution. Most Dm types have higher coverage factors (Fig. 5c), so that breaking of translation invariance is mild. But Dm4 has a low coverage factor (Fig. 5a, S4), so translation invariance of its target Mi4 would be broken quite severely (Fig. 5b, c) if lateral inhibition were the function of Dm4. If, on the other hand, Dm4 terminals function as independent units, then perfect tiling of Dm4 would not break translation invariance. Dm4 would be much like an assembly of unicolumnar cells, implementing convolution with stride one.

The higher coverage factors of most Dm types (Fig. 5d) are also typical of most ganglion cell types in the mouse retina (Bae et al. 2018). A higher coverage factor means that translation invariance is approximately preserved, which may be a design principle in visual systems.

### Dm4 may still act as a functional unit for top-down inputs

If the columns of a Dm4 arbor are electrically independent, could there be any advantage of uniting them into a single cell? One intriguing possibility is that the columns are not electrically independent when the cell receives synapses from centrifugal pathways originating in the central brain. Such synapses tend to lie on thicker dendrites that are closer to the cell body than the terminals (data not shown). In a passive dendrite, electrical attenuation along the thick-to-thin direction is less than along the thin-to-thick direction (Rall and Rinzel 1973; Roth and Häusser 2001). Therefore, synaptic input on the thicker portions of dendrites may propagate to the Dm4 terminals with relatively little attenuation, so that top-down input influences all terminals as one functional unit.

### Comparison with physiology

The structural analysis of Fig. 2 can be compared with physiological measurements of the Mi4 receptive field. The sizes of the ON center and OFF surround of the Mi4 receptive field as measured by white noise analysis (Arenz et al. 2017) roughly match the direct pathway L5-Mi4 (Fig. 2d) and the L5-Dm10-Mi4 inhibitory field (Fig. 2e), respectively.

The L2-Dm15-L2 inhibitory field is anisotropic, with a horizontal orientation (Fig. 2g). This is reminiscent of the old observation of a horizontally oriented inhibitory field in the *Limulus* eye (Barlow 1969). The L2-Dm15-L2 inhibitory field is expected to give rise to a horizontally oriented surround for the L2 receptive field. Such an anisotropic surround has not been reported in experiments with white noise stimuli (Drews et al. 2020). The effect might be subtle, as L2 receives input from many sources (Fig. S3) and the L2 receptive field is likely created by multiple overlapping mechanisms. The role of GABAergic inhibition in creating the surround of the L2 receptive field was previously investigated (Freifeld et al. 2013). Knockdown of GABA receptors in L2 cells alone had no effect on the receptive field. This is not inconsistent with the idea that Dm15 contributes to the L2 surround, because Dm15 is predicted glutamatergic.

Contrast normalization has been studied by calcium imaging of the columnar types of Fig. 1a, and the observations were fit by neural network models (Drews et al. 2020). Luminance normalization was studied by calcium imaging of Tm1 and Tm9, and modeled with a neural network (Gür et al. 2024). A proposed mechanism for normalization was that Dm12 pools L3 inputs and inhibits Tm9. In Fig. 3a, Tm9 is suppressed in the top motif because it is a relatively weak target of Dm12, but it is a target nonetheless. Adding it to Fig. 3a would result in a combination of recurrent and feedforward inhibition similar to the motifs of Fig. 3d. According to this combined motif, strong recurrent inhibition of L3 by Dm12 is also expected to normalize L3 input to Tm9. Disentangling the effects of these parallel mechanisms on Tm9 responses will require an interplay between detailed physiology and neural network modeling.

### Is lateral inhibition subtractive or divisive?

The connectivity motifs of this paper will be important for constraining network models of normalization (Drews et al. 2020; Gür et al. 2024). One important issue for all such models is whether inhibition is subtractive or divisive (Furman 1965). In the subtractive model, the currents from excitatory and inhibitory synapses are approximated as combining linearly as in Kirchhoff’s First Law. The divisive or shunting model arises from the ratio of current and conductance in Ohm’s Law, because synapses affect voltage by causing conductance changes. The center-surround receptive field in the *Limulus* eye was modeled with subtractive inhibition by (Hartline and Ratliff 1958). The model of human spatial vision due to (Sperling 1970) is divisive.

Normalization in convolutional nets combines subtractive and divisive steps (Ioffe and Szegedy 2015; Ulyanov, Vedaldi, and Lempitsky 2016). The biologically inspired forerunners of convolutional nets made use of divisive normalization (Fukushima 1975, 1980), as did influential models of mammalian neurophysiology (Carandini and Heeger 1994, 2011).

The models mentioned above have assumed that inhibition is divisive (Drews et al. 2020; Gür et al. 2024). There is also direct electrophysiological evidence that inhibition is divisive in fly T4 visual neurons (Groschner et al. 2022). Physiological studies of inhibition from Dm interneurons are clearly warranted.

Whether inhibition is divisive or subtractive has been controversial for mammalian neurons (Holt and Koch 1997; Silver 2010). Many theoretical studies have attempted to relate this question to synapse location, motivated by the fact that mammalian interneuron types often prefer to synapse on particular compartments of the postsynaptic neuron. In the fly medulla, Dm neurons synapse onto the axons of L cells and onto the dendrites of other columnar types, and these facts might be relevant for understanding the effects of inhibition.

### Luminance vs. contrast encoding

From the above, it is clear that modeling normalization in the fly optic lobe will require careful consideration of circuit connectivity as well as cellular biophysics, and is outside the scope of the present work. Here I venture only some simple remarks about this complex subject.

Since L2 receives three types of Dm-mediated lateral inhibition (Fig. 4a), its output is expected to be highly normalized. In contrast, L1 has little or no connectivity with Dm types (Fig. S1, S2), so its output is expected to be relatively unnormalized. L3 receives lateral inhibition in the lamina (Fig. 3g), and its lateral inhibition from Dm12 (Fig. 3a) is rather short-range (Fig. 4g). Therefore L3 is expected to be less normalized than L2 but more than L1.

L2 is insensitive to luminance (Ketkar et al. 2020), consistent with L2 being highly normalized. L1 (Ketkar et al. 2022) encodes luminance rather linearly, consistent with the idea of no normalizing mechanisms in the medulla. L3 (Ketkar et al. 2020) encodes luminance but rather nonlinearly, consistent with some degree of normalization.

These arguments are merely hand-waving preludes to future computational models of normalization in fly vision. They serve to illustrate the general idea that the connectivity of Dm types should have strong implications for visual response properties of columnar neurons.

### Diversity and specificity

In the fly optic lobe, the reason for a large number of interneuron types may be simple. Diversity is an inevitable consequence of specificity. This is also the case in convolutional nets, which contain a large number of highly specific normalizations. The relevance of these ideas to mammalian cortex is currently unclear. There is some physiological evidence that normalization is specific to particular visual features such as orientation (Self et al. 2014; Trott and Born 2015). However, current models of cortical normalization invoke only one or a few types of interneuron (Obeid and Miller 2021; Heeger and Zemlianova 2020), far less than the 60 interneuron types suggested by transcriptomic studies of mouse visual cortex (Tasic et al. 2018).

### Functional annotation of interneuron types

In addition to analyzing Dm circuits in detail, I also searched for candidate normalizers in other interneuron families throughout the optic lobe (Fig. 6). Because the relevant motifs are so widespread, a significant fraction of optic lobe cell types seem likely to be normalizers. This shows the power of connectomics to provide clues to the functions of cell types before any visual physiology experiments have been performed.

The present work can be regarded as a contribution to the emerging field of connectome annotation (Scheffer et al. 2020; Dorkenwald et al. 2023; Schlegel et al. 2023). Connectomic analysis provides not only cell types but also rules of their connectivity. These rules can be used to deduce functional annotations for the cell types, and the present work shows by example how to arrive at such deductions. The traditional method of investigating the function of a visual neuron has been to record its responses to visual stimuli, but such experiments are time-consuming and laborious. I expect that such physiological studies will be accelerated by functional annotations of the connectome like the ones provided here.

## Methods

All analyses are based on v783 of the FlyWire neuronal wiring diagram of a female adult fly brain (Zheng et al. 2018; Dorkenwald et al. 2023; Schlegel et al. 2023), and cell types assigned in the right optic lobe (Matsliah et al. 2023). Cells were assigned to a hexagonal lattice of columns as described in a companion paper (Seung 2023).

### Input and output fractions

Let *N*_A→B_ be the number of synapses from type A neurons to type B neurons. This number is normalized to yield the input fraction,

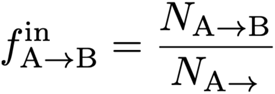

Here *N*_A→_ is the number of synapses from A neurons to any neuron, and *N*_→B_ is the number of synapses from any neuron to B neurons. The output fraction is analogously defined as

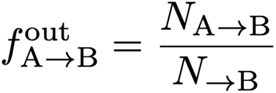

## Acknowledgements

I thank Ben Silverman for the Dm4 skeleton tracing of Fig. 5d, Ryan Willie for illustrations in Fig. 1, and the FlyWire team for the connectome and cell types on which this analysis is based. I am grateful to Michael Hausser for advice concerning dendritic biophysics, to Ken Miller for information about cortical interneurons, and to Yijie Yin for corrections to the manuscript.

## Supplementary Figures

**Figure S1.**
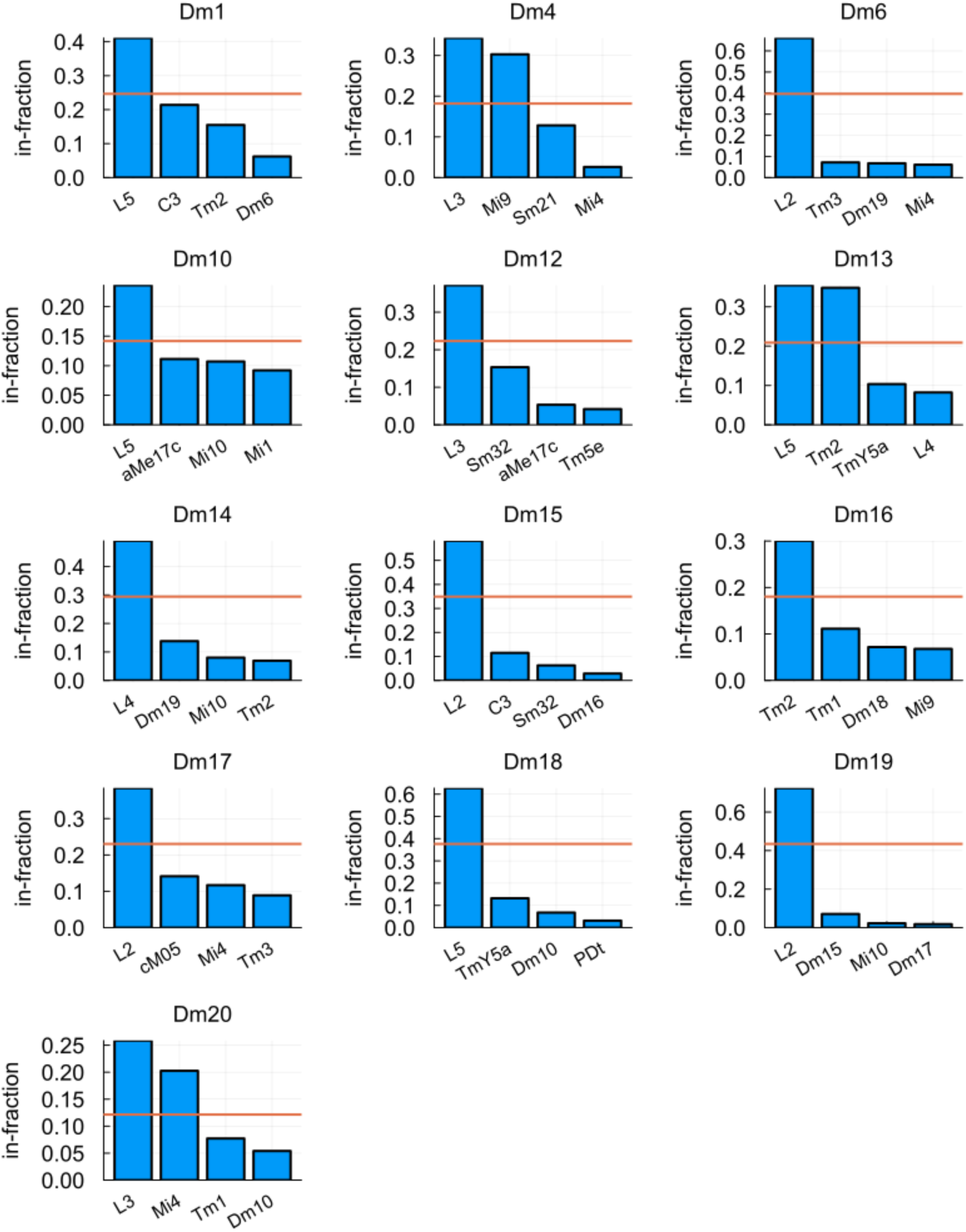
Each Dm type is dominated by one or two input types. Top four input types for each Dm type. The bars poking above the horizontal red line are said to be dominant inputs. The red line is placed at 60% of the height of the shortest bar above the red line. Most Dm types have a single dominant input. Dm13, Dm20, and Dm4 have two dominant inputs.

**Figure S2.**
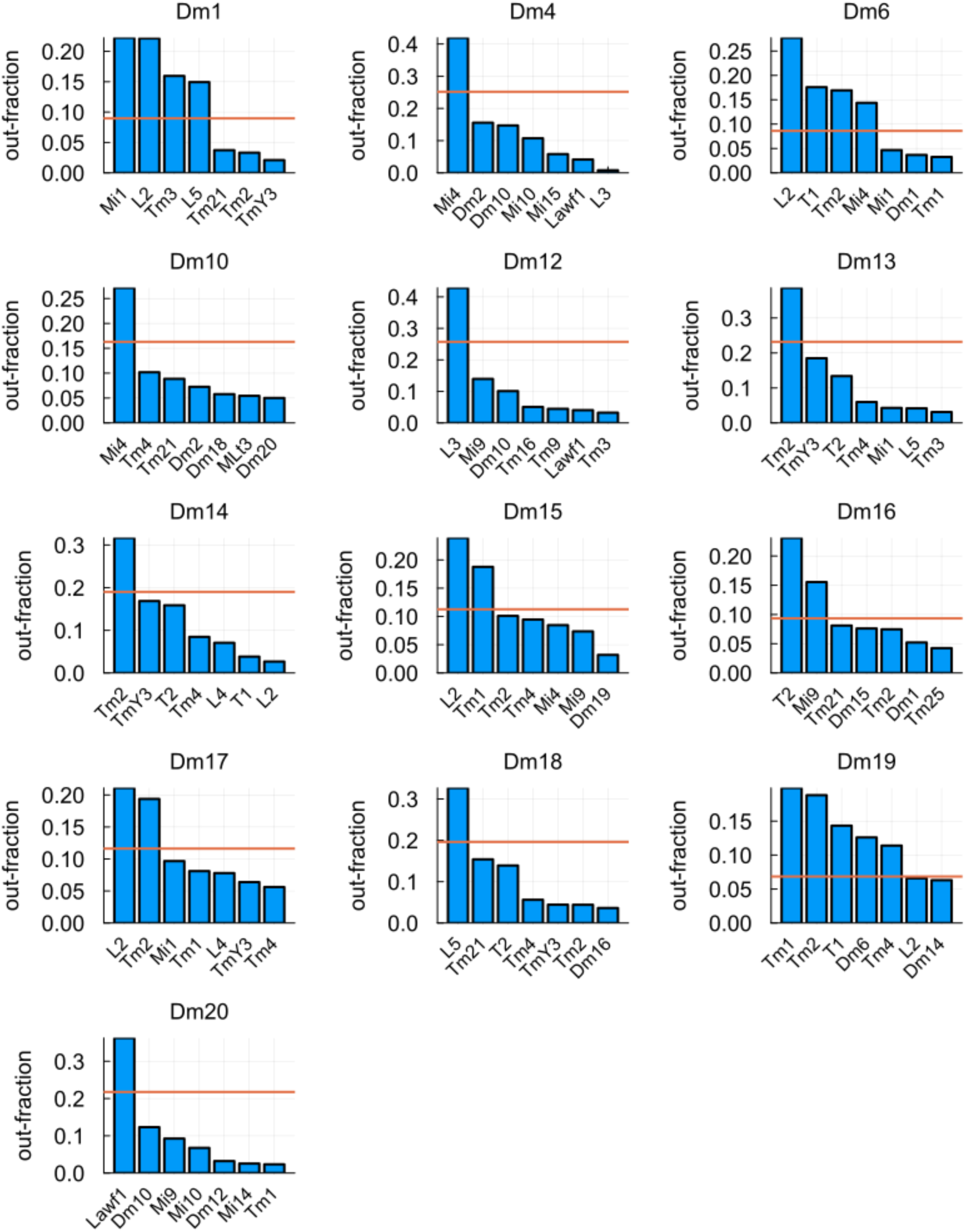
Dominant outputs for Dm interneurons. Top six output types for each Dm type. The bars poking above the horizontal red line are said to be dominant outputs. The red line is placed at 60% of the height of the shortest bar above the red line. Roughly half the Dm types have a single dominant output, but some have four or even five.

**Figure S3.**
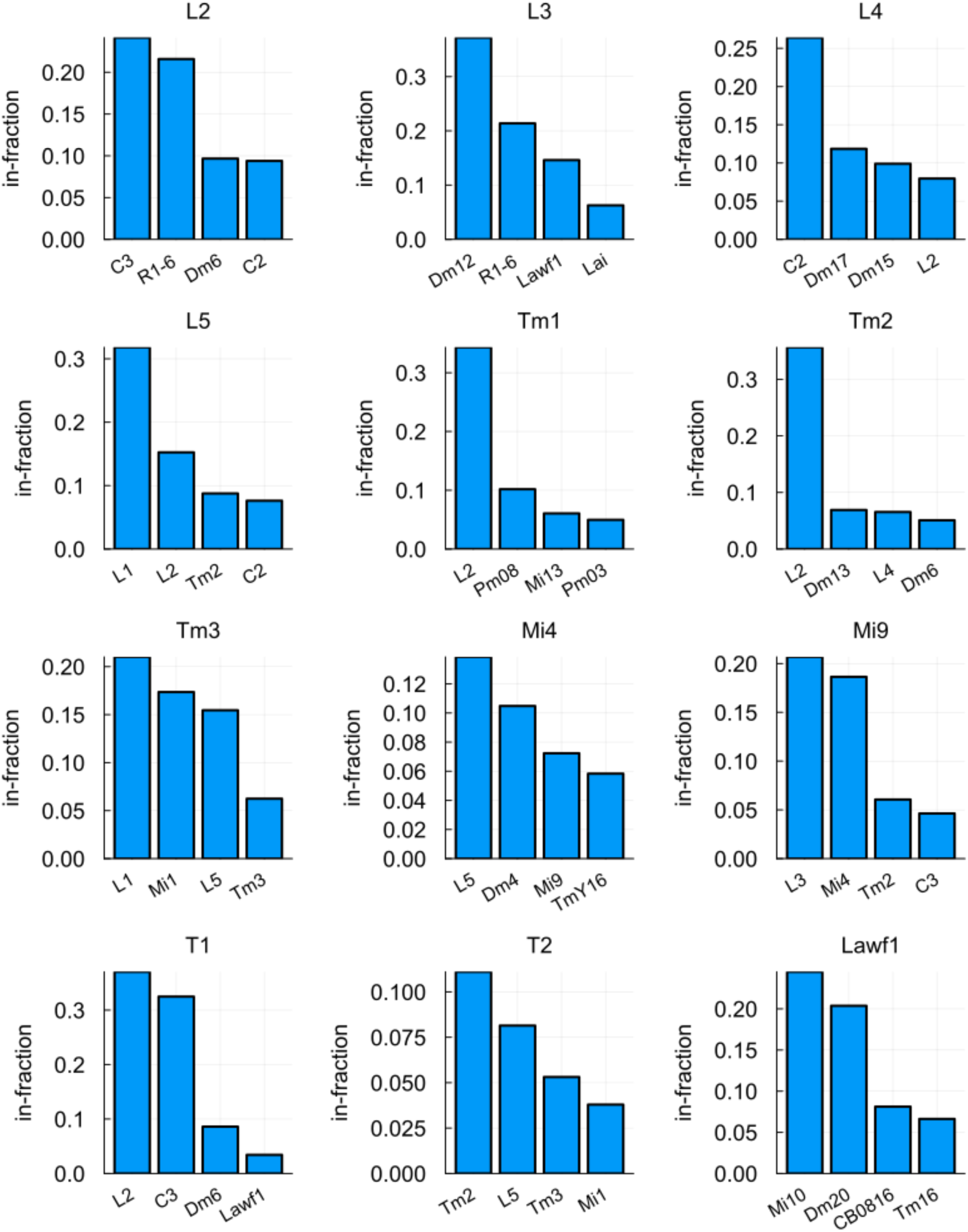
Inputs to columnar types. Top four inputs to the columnar types that are the main partners (dominant inputs or outputs) of Dm types (same columnar types as Fig. 1e). R1-6 input fractions are known to be inaccurate due to underdetection of photoreceptor synapses, and a fix is in progress.

**Figure S4.**
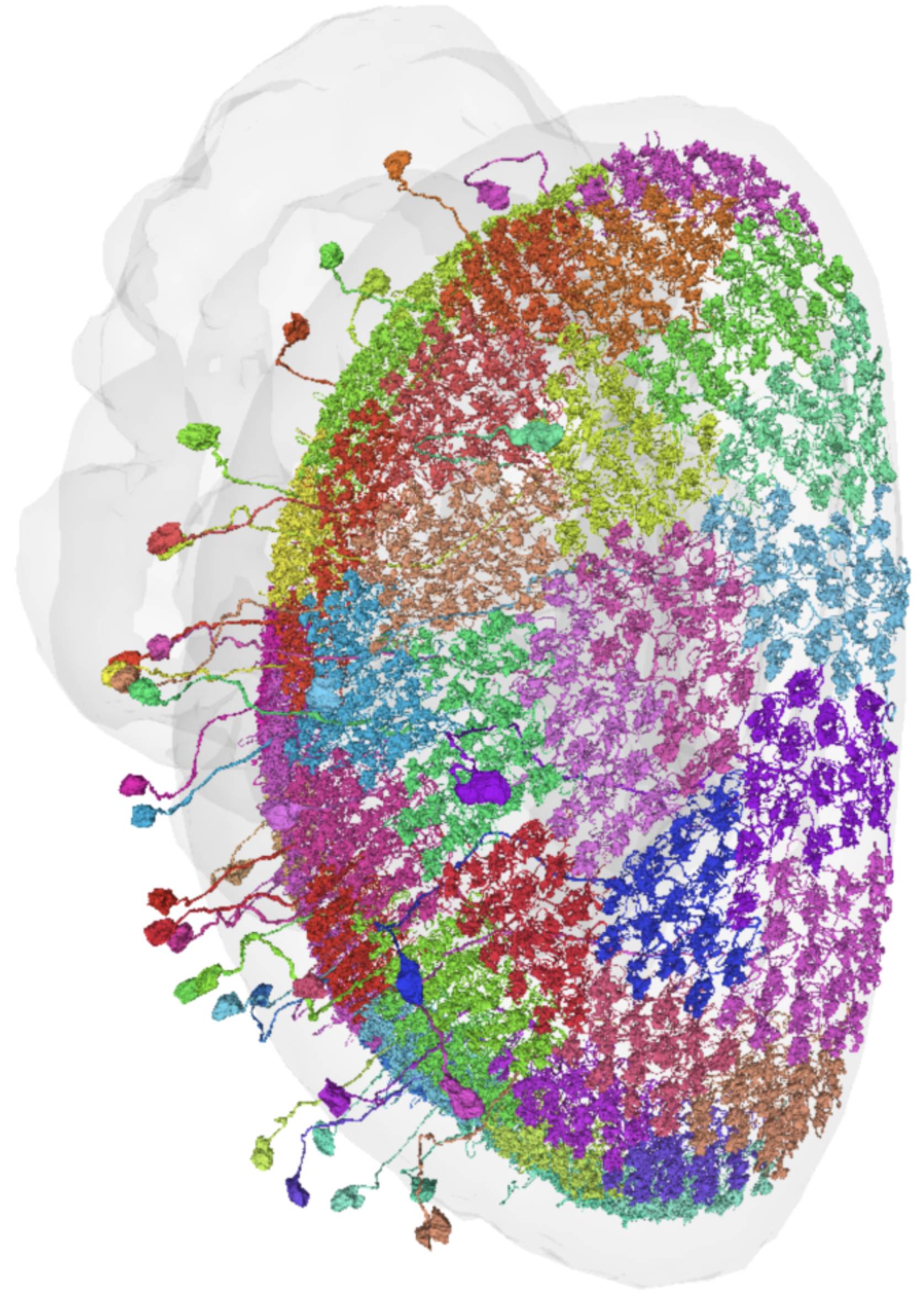
Perfect tiling of the visual field by Dm4 cells. Dm4 is the only type in the Dm4 baker’s dozen that perfectly tiles the medulla, leading to a coverage factor that is approximately one (Fig. 5a). Other Dm types contain cells that overlap significantly with each other, with coverage factors that are typically 2 or 3 (Fig. 5a).

**Figure S5.**
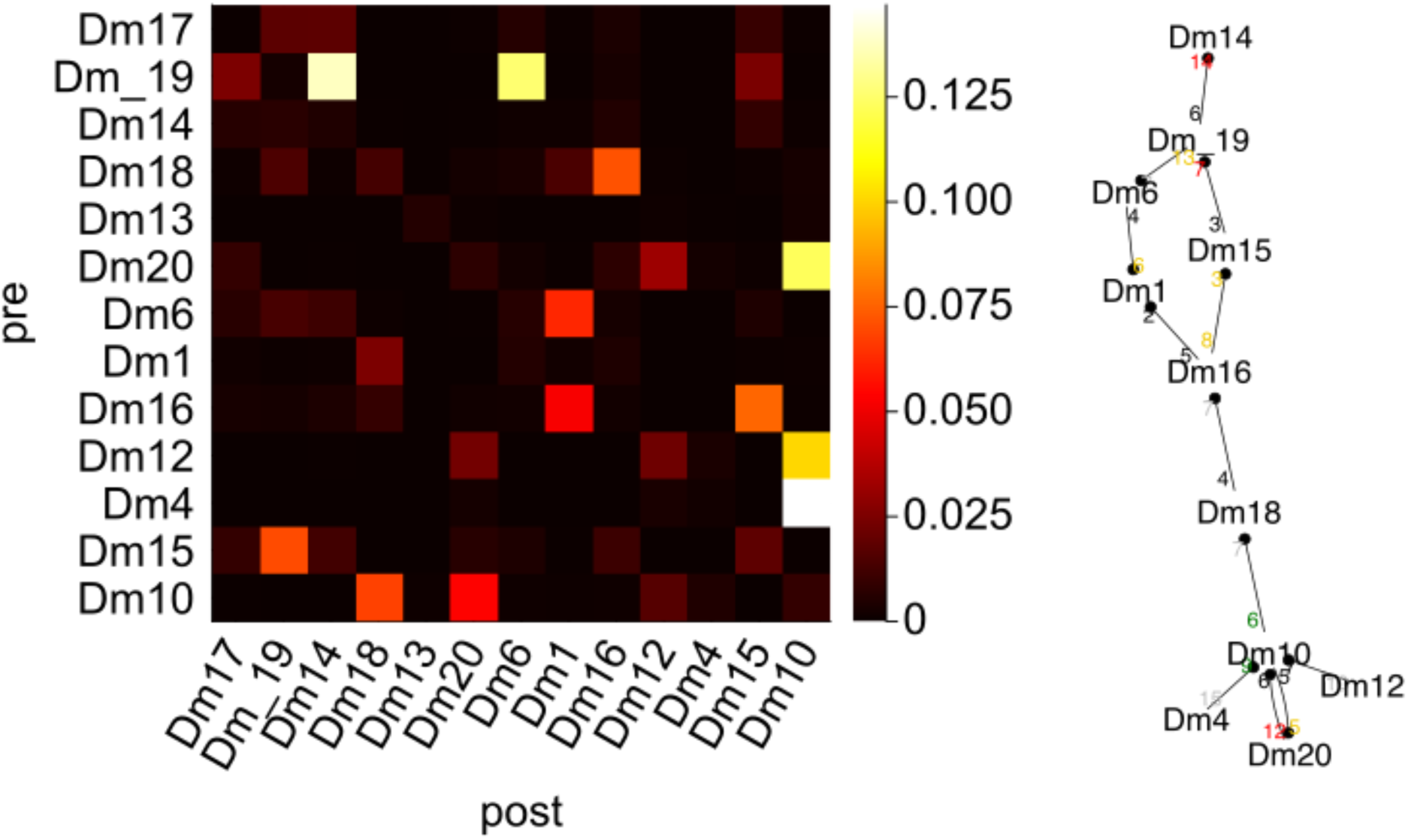
Dm-Dm disinhibition. (a) Connectivity between Dm types, ordered with decreasing arbor size. Each pixel value denotes “importance,” defined as the maximum of the input and output fractions for that connection. Pixels above the diagonal correspond to connections from larger to smaller cells, and vice versa for pixels below the diagonal. (b) Graph showing connections with importance greater than 5%.

## Footnotes

1 Dm12 is a recent exception, as visual responses have now reported in a preprint (Gür et al. 2024).

